# A highly conserved host lipase deacylates oxidized phospholipids and ameliorates acute lung injury

**DOI:** 10.1101/2021.07.26.453786

**Authors:** Benkun Zou, Michael Goodwin, Danial Saleem, Wei Jiang, Jianguo Tang, Yiwei Chu, Robert Munford, Mingfang Lu

## Abstract

Oxidized phospholipids have diverse biological activities, many of which can be pathological, yet how they are inactivated *in vivo* is not fully understood. Here we present evidence that a previously unsuspected host lipase, acyloxyacyl hydrolase (AOAH), can play a significant role in reducing the pro-inflammatory activities of two prominent products of phospholipid oxidation, 1-palmitoyl-2-glutaryl-sn-glycero-3- phosphocholine (PGPC) and 1-palmitoyl-2-(5-oxovaleroyl)-sn-glycero-3-phosphocholine (POVPC). AOAH removed the sn-2 and sn-1 acyl chains from both lipids and reduced their ability to induce macrophage inflammasome activation and cell death *in vitro* and acute lung injury *in vivo*. In addition to transforming Gram-negative bacterial lipopolysaccharide from stimulus to inhibitor, its most studied activity, AOAH can inactivate these important danger-associated molecular pattern (DAMP) molecules and reduce tissue inflammation and cell death.

## Introduction

Oxidized phospholipids (oxPLs) are produced by peroxidation of cell membrane or lipoprotein phospholipids, usually during inflammation or senescence (Bochkov et al., 2010). They can be recognized by many receptors in the innate immune system, often with significant consequences (Bochkov et al., 2017; Bochkov et al., 2010; Erridge et al., 2008; Hazen, 2008; Matt et al., 2015; Mauerhofer et al., 2016). Some oxPLs may induce cell death or endothelial barrier disruption (Fruhwirth and Hermetter, 2008). For example, 1-palmitoyl-2-glutaroyl-sn-glycero-3-phosphocholine (PGPC) and 1-palmitoyl- 2-(5-oxovaleroyl)-sn-glycero-3-phosphocholine (POVPC), two important oxidation products of 1-palmitoyl-2-arachidonoyl-sn-glycero-3-phosphocholine (PAPC), can activate acid sphingomyelinase to produce ceramide that then induces cell death of smooth muscle cells and macrophages (Greig et al., 2012; Loidl et al., 2003; Stemmer et al., 2012). In endoplasmic reticulum-stressed macrophages, oxPLs may also trigger apoptosis in a CD36-TLR2 dependent manner (Seimon et al., 2010), and ozone-treated lung surfactant component 1-palmitoyl-2-(9’-oxo-nonanoyl)-sn-glycero-3- phosphocholine can induce macrophage death (Uhlson et al., 2002). OxPAPC components such as PGPC and POVPC may disrupt endothelial barriers in the lung while a full-length oxygenated product, 1-palmitoyl-2-(5,6-epoxyisoprostane E2)-sn-glycero-3- phosphatidyl choline (PEIPC), was found to protect the barrier (Birukova et al., 2013).

In addition to inducing cell death, oxPLs are able to evoke inflammatory responses in macrophages (Bochkov et al., 2010; Vladykovskaya et al., 2011). For example, PGPC and POVPC may induce inflammasome activation and IL-1β release from macrophages that have been primed by LPS or other TLR agonists (Yeon et al., 2017; Zanoni et al., 2017). Unlike other stimulus-induced inflammasome activators, however, oxPLs can induce IL-1β release without inducing pyroptosis (Evavold et al., 2018; Hagar et al., 2013; Kayagaki et al., 2011; Kayagaki et al., 2013; Mangan et al., 2018; Shi et al., 2015; Shi et al., 2014; Zanoni et al., 2016; Zanoni et al., 2017). Building on evidence that cell- surface CD14 can bind and internalize glycerophosphoinositides (Wang et al., 1998; Wang and Munford, 1999), Zanoni et al. found that CD14 captures and internalizes PGPC and POVPC and showed that this mode of cell entry is required for these oxPLs to induce macrophages to release IL-1β (Zanoni et al., 2017). Although macrophages play an essential role in many inflammatory responses, how they degrade oxPLs and the biological impact of this degradation are not well understood (Zanoni et al., 2017).

Acyloxyacyl hydrolase (AOAH), a highly conserved animal lipase, is expressed in phagocytes, including macrophages, microglia, dendritic cells and neutrophils, and also in NK cells, ILC1s and renal proximal tubule cells (Munford et al., 2020). AOAH transforms bacterial lipopolysaccharide (LPS) by cleaving the ester bonds that attach the secondary (“piggyback”) fatty acyl chains to the primary (glucosamine-bound) hydroxyacyl chains in the lipid A moiety, converting a potent stimulus to a competitive inhibitor (Munford et al., 2020). In addition, AOAH can remove fatty acyl chains from the sn-1 and sn-2 positions of many glycero (phospho) lipids and be an acyl transferase (Gorelik et al., 2018; Munford and Hunter, 1992; Staab et al., 1994).

Here we present evidence that AOAH can (a) reduce the ability of two prominent oxPLs (PGPC and POVPC) as well as an intermediate product of their deacylation (lysophosphatidycholine, LPC) to induce cell death and IL-1β release from macrophages *in vitro* and (b) prevent oxPL-induced acute lung injury *in vivo* in mice.

## Results

### Low concentrations of oxPLs induce cell death and IL-1β release from primed macrophages

We began by asking if low concentrations of oxPAPC, PGPC and POVPC can induce cell death and IL-1β release from mouse resident peritoneal macrophages. Cells were either not primed or primed with 10 ng/ml TLR2/1 agonist Pam3CSK4 (Pam3) for 7 h and then a test lipid (PGPC, POVPC, oxPAPC, or non-oxidized PAPC) was added to the culture media for an additional 18 h. We found that 5 – 12.5 μg/ml (8 – 21 μM) PGPC or POVPC and 12.5 μg/ml oxPAPC could induce significant cell death with or without Pam3 priming, whereas 12.5 μg/ml (16 μM) non-oxidized PAPC was inactive (Fig. 1A). PGPC- and POVPC-induced IL-1β secretion occurred only in Pam3-primed cells and was dose-dependent (Fig. 1B). Non-oxidized PAPC did not induce IL-1β release by primed cells (Fig. 1B), in agreement with previous findings that only oxidized PLs activate the inflammasome (Zanoni et al., 2016; Zanoni et al., 2017). Pam3-induced IL-6 production was minimally affected by the presence of oxPLs (Fig. 1C). Mature IL-1β (p17) and cleaved caspase 1 (p20) appeared in the culture medium only after Pam3-primed macrophages had been treated with either PGPC or POVPC (Fig. 1D, E). Caspase 1 inhibitor VX-765 and pan-caspase inhibitor Z-VAD-FMK significantly reduced IL-1β and cleaved caspase 1 (p20) release in primed macrophages but did not prevent oxPL- induced cell death, demonstrating that oxPL-induced cell death is independent of inflammasome activation (Stemmer et al., 2012) (Fig. S1A-C and Fig. 1E). In addition, we found that caspase 11 is not required for PGPC-induced IL-1β release from primed peritoneal macrophages (Fig. S1, D-F) (Zanoni et al., 2016). Low concentrations of oxPLs thus induced cell death; inflammasome-produced IL-1β and cleaved caspase 1 were released if the macrophages had been primed.

**Fig. 1.**
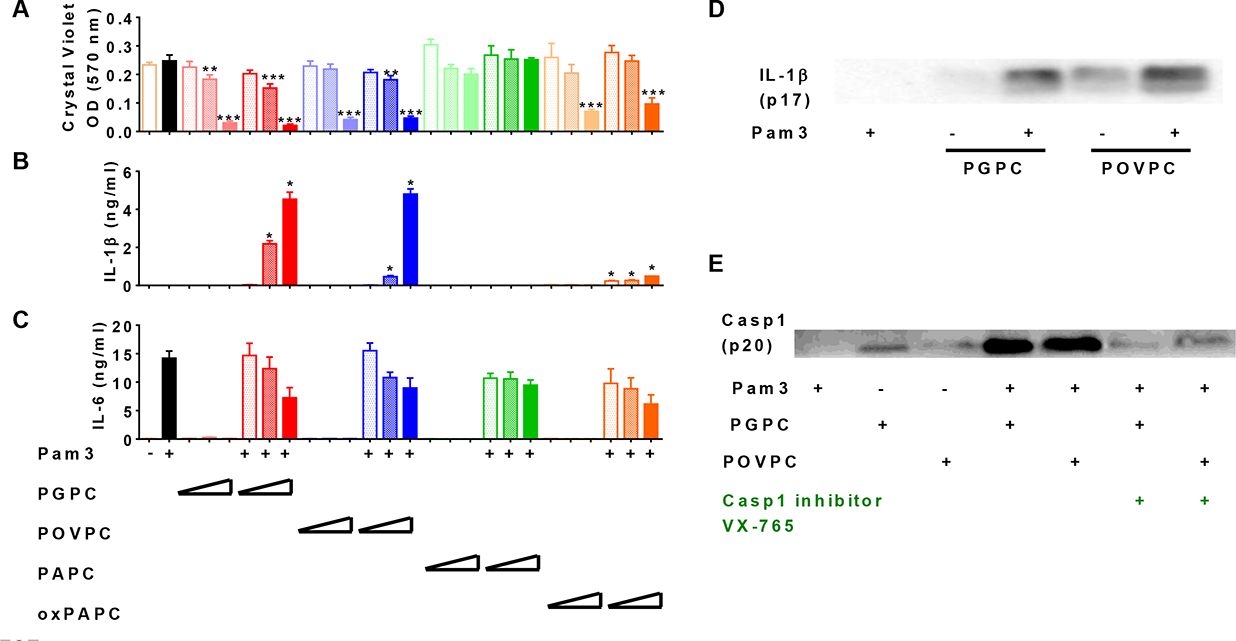
Low concentrations of oxPLs induce macrophage cell death but IL-1β release requires priming. **(A - C)** Resident peritoneal macrophages were incubated for 7 h with or without priming with 10 ng/ml Pam3. Two, 5, or 12.5 μg/ml PGPC, POVPC, PAPC or oxPAPC were added to the media. After incubation for 18 h, cells were washed and then stained with crystal violet (A). IL-1β and IL-6 were measured in the culture media using ELISA (B, C). One representative experiment of 4 is shown, n = 3 or 4 wells/group/experiment. **(D, E)** Macrophages were treated with 10 ng/ml Pam3 for 7 h and then 12.5 μg/ml PGPC or POVPC was added. After 18 h incubation, the media were concentrated and subjected to Western blot analysis for mature IL-1β (D) and cleaved caspase 1 (p20) (E).

### AOAH deacylates and inactivates oxPLs

AOAH can act *in vitro* as a phospholipase A_1_, A_2_ or B (Gorelik et al., 2018; Munford and Hunter, 1992). To find out if the enzyme can deacylate oxPLs, we incubated PGPC and POVPC with purified recombinant human AOAH (rhAOAH) for 2 or 4 h. Analysis using LC-MS showed that AOAH can remove the sn-2 oxidized fatty acyl chain and sn-1 palmitate from both PGPC and POVPC (Fig. 2A, B); the sn-2 moiety was removed preferentially, in keeping with the enzyme’s preference for removing shorter acyl chains from glycerolipids (Erwin and Munford, 1990; Gorelik et al., 2018; Munford and Hunter, 1992). AOAH treatment greatly reduced the ability of PGPC and POVPC to induce cell death (Fig. 2C, D) and IL-1β release (Fig. 2E, F) in Pam3-primed macrophages, while Pam3 stimulated similar levels of IL-6 production (Fig. 2E) in the presence of either oxPLs or deacylated oxPLs. AOAH thus decreased the ability of these oxPLs to induce cell death and inflammasome activation.

**Fig. 2.**
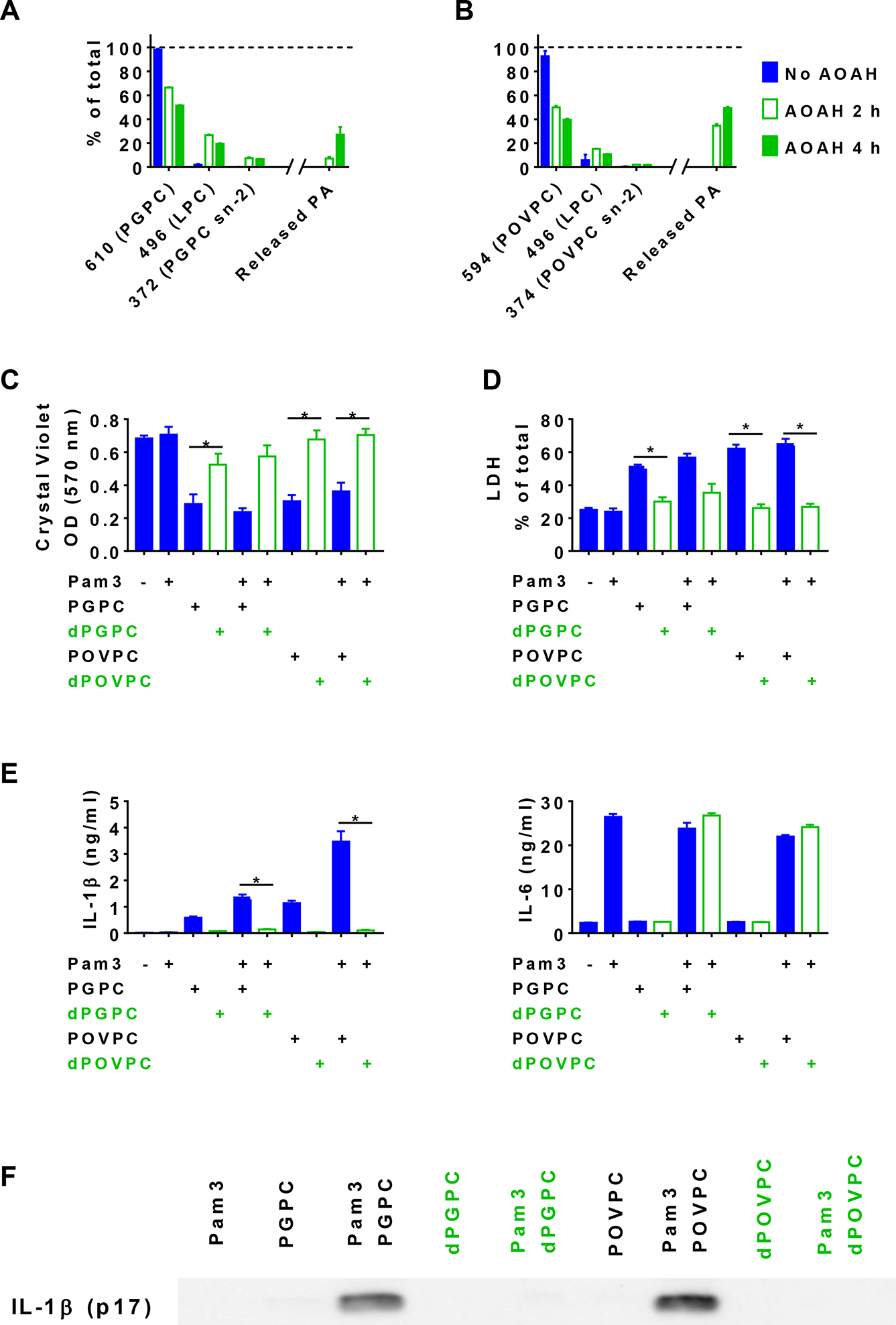
AOAH treatment decreases the bioactivities of PGPC and POVPC. **(A, B)** PGPC and POVPC were incubated with rhAOAH at 37℃ for 2 or 4 h in 100 mM NaCl with 10 mM sodium acetate, pH 6.2, 0.1% Triton X-100, and 0.2 mg/ml fatty acid-free human BSA. The reaction products missing sn-2 oxidized FA (MW 496, LPC) or missing sn-1 palmitic acid (MW 372 for PGPC or MW 374 for POVPC) were analyzed (see Methods); the released palmitic acid (PA) was also measured. n = 3 or 4. **(C - G)** Resident peritoneal macrophages were treated or untreated with 10 ng/ml Pam3 for 7h. PGPC, POVPC or AOAH-deacylated PGPC or POVPC (dPGPC or dPOVPC) (12.5 μg/ml) was then added. After incubation for 18 h, cells were stained with crystal violet and the OD was measured (C). Released LDH was measured (D). n = 4. Concentrations of IL-1β and IL-6 (E) were measured in the culture media, which were also subjected to Western blot analysis (F). For the data in C – F, one representative experiment of 3 or 4 is shown.

### AOAH also deacylates and inactivates LPC

When AOAH released the sn-2 oxidized acyl chains from PGPC or POVPC, the major product was LPC (Fig. 2A, B), another bioactive DAMP with disease associations (Liu et al., 2020); the appearance of free palmitate was evidence for AOAH-mediated deacylation of LPC (Munford and Hunter, 1992). When we incubated pure LPC with rhAOAH we again found that palmitate was released (Fig. 3A). In agreement with previous reports that LPC can be inflammatory and induce IL-1β release (Li et al., 2016; Liu-Wu et al., 1998; Vladykovskaya et al., 2011), we found that LPC induced cell death and IL-1β release by macrophages; AOAH treatment partially diminished these bioactivities, including a major reduction in IL-1β release (Fig. 3B-F). In response to LPC, Pam3-primed *Aoah^-/-^* macrophages released more caspase 1 p20 than did primed *Aoah^+/+^* macrophages (Fig. 3G). Thus, AOAH can deacylate and inactivate LPC *in vitro* and prevent LPC-induced inflammasome activation.

**Fig. 3.**
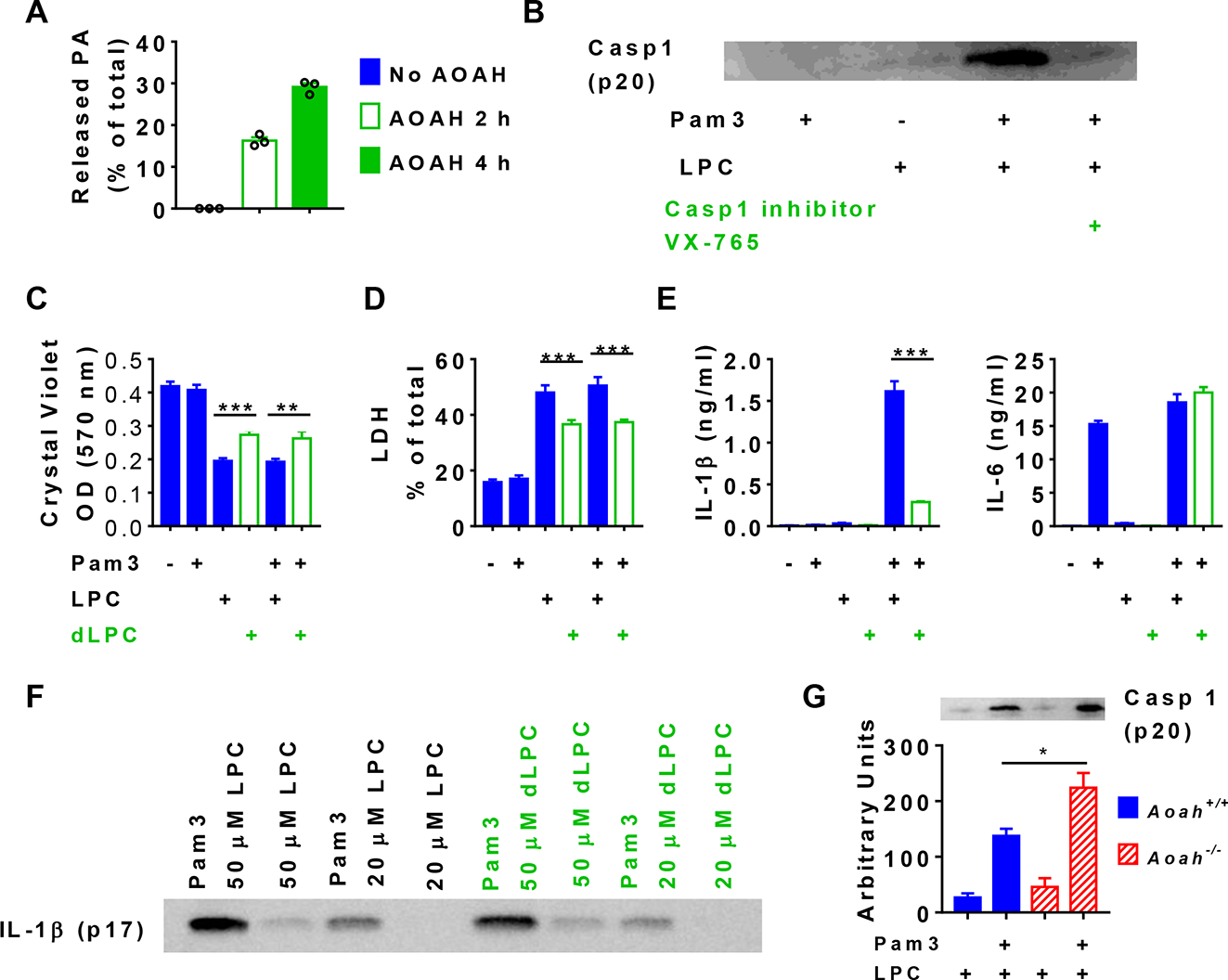
AOAH treatment decreases the bioactivity of LPC. **(A)** After LPC was incubated with purified rhAOAH for 2 or 4 h, the released palmitate was measured. No palmitate was detected when rhAOAH was absent. n = 3. **(B)** Resident peritoneal macrophages were incubated for 7 h with or without priming with 10 ng/ml Pam3 before 12.5 μg/ml (25 μM) LPC was added to the media. After 18 h incubation, cleaved caspase 1 (p20) in the culture medium was analyzed by Western blot. Ten μM VX-765 was added 1 h before Pam3 treatment in some wells. **(C – F)** Resident peritoneal macrophages were treated or untreated with 10 ng/ml Pam3 for 7 h before 12.5 μg/ml (25 μM) LPC or AOAH deacylated LPC (dLPC) was added. After incubation for 18 h, cells were stained with crystal violet and the media was used to measure LDH activity (C), IL-1β and IL-6 concentrations (D, E), and mature IL-1β (Western blot) (F). (C - E) Data were combined from 2 or 4 experiments, each with n = 4. **(G)** *Aoah^+/+^*and *Aoah^-/-^* macrophages were treated with 10 ng/ml Pam3 for 7 h and then 5 μg/ml LPC was added. After 18 h incubation, the media were subjected to Western blot analysis for cleaved caspase 1 (p20). Western blot results from 3 experiments were quantitated using Image J and plotted with the Western results. n = 5 wells/group.

### AOAH reduces oxPL-induced cell death and inflammasome activation in macrophages

We next explored a potential role for AOAH in inactivating oxPL in cells by studying the responses of *Aoah^+/+^* and *Aoah^-/-^* resident peritoneal macrophages to PGPC and POVPC. With or without Pam3 priming, PGPC or POVPC treatment led to more cell death in *Aoah^-/-^* macrophages than in *Aoah^+/+^* controls (Fig. 4A). After Pam3-priming, oxPLs induced more IL-1β release from *Aoah^-/-^* macrophages than from *Aoah^+/+^* cells (Fig. 4B). Although AOAH can deacylate synthetic lipopeptides, including Pam3 (Gorelik et al., 2018), AOAH did not regulate IL-6 secretion in response to Pam3 and/or oxPL stimulation (Fig. 4C). Therefore, AOAH does not prevent macrophages from responding to Pam3, which can initiate cell signaling by engaging cell-surface TLR2, in keeping with previous findings that AOAH does not modulate acute responses to LPS, such as secretion of TNF-α, IL-6 or RANTES (Lu et al., 2008). In addition, we found that oxPLs induced significantly greater release of cleaved caspase 1 in Pam3-primed *Aoah^-/-^* macrophages, confirming that AOAH diminishes oxPL-induced inflammasome activation (Fig. 4D). When oxPLs are added to culture media, CD14-dependent internalization of oxPLs is required for inflammasome activation (Zanoni et al., 2017). When we delivered PGPC or POVPC into the cell cytosol using DOTAP, bypassing CD14 and endosomes (Zanoni et al., 2017), primed *Aoah^+/+^* and *Aoah^-/-^* macrophages released similar amounts of cleaved caspase 1 (p20) (Fig. 4E). In addition, AOAH did not alter cytosolic LPS-induced IL-1β release (Fig. 4F) (Shi et al., 2014; Zanoni et al., 2016) or modulate ATP-induced inflammasome activation or IL-1β release in Pam3-primed macrophages (Fig. 4G), evidence that AOAH does not prevent inflammasome activation *per se*. These results suggest that oxPL internalization via CD14-induced endocytosis may allow AOAH, which is found in endo-lysosomes (Luchi and Munford, 1993; Staab et al., 1994), to deacylate oxPLs before they activate the inflammasome (Zanoni et al., 2017).

**Fig. 4.**
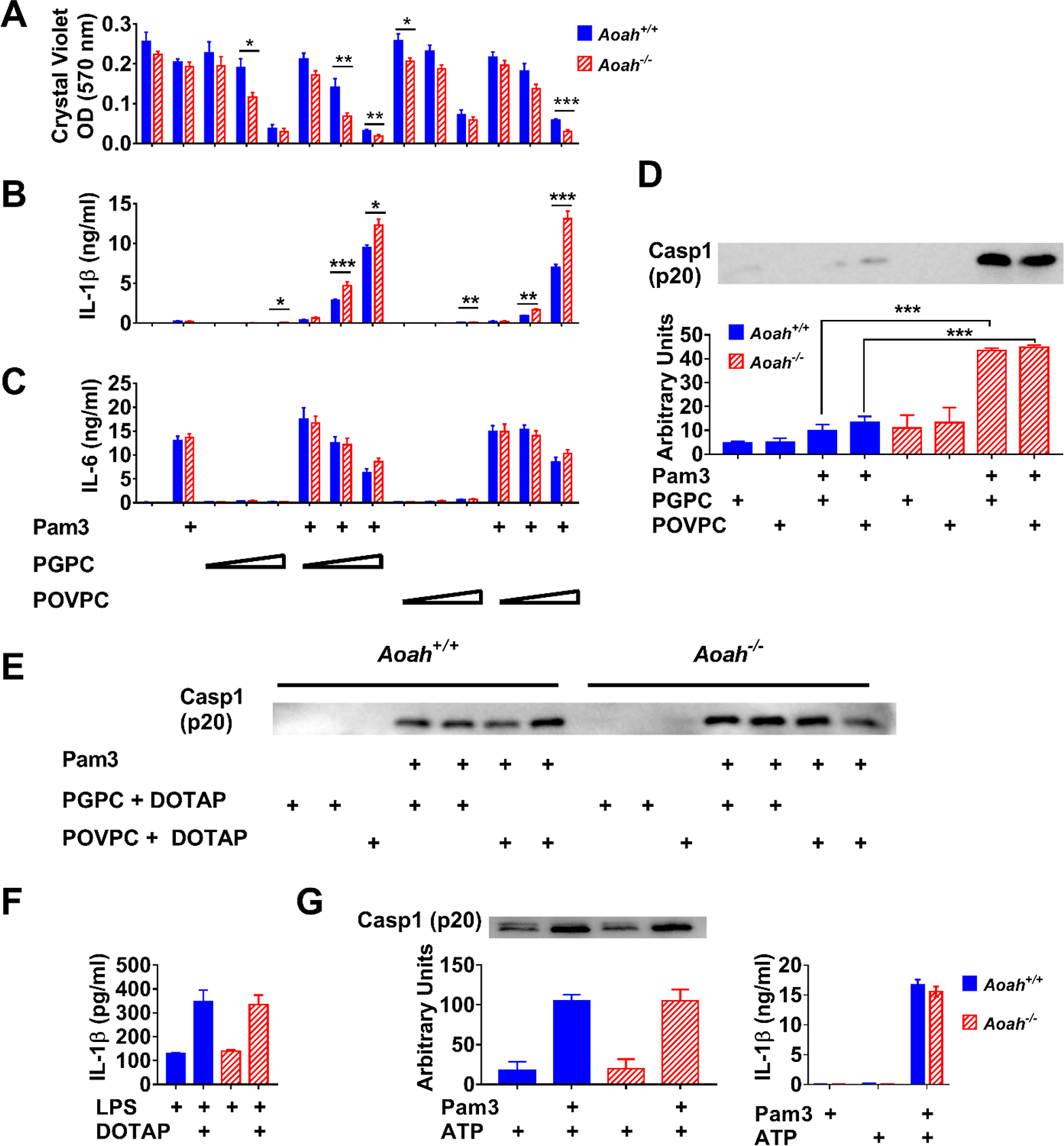
AOAH reduces oxPL-induced cell death and inflammasome activation in macrophages. **(A - C)** Resident peritoneal macrophages were incubated for 7 h with or without priming with 10 ng/ml Pam3. Two, 5, or 12.5 μg/ml PGPC or POVPC were then added to the media. After incubation for 18 h, cells were washed and then stained with crystal violet (A). IL-1β (B) and IL-6 (C) were measured in the culture media using ELISA. Data were combined from 3 experiments, each with n = 4 wells/group. **(D)** Macrophages were primed with 10 ng/ml Pam3 and then 5 μg/ml PGPC or POVPC was added. After 18 h incubation, the media were subjected to Western blot analysis for cleaved caspase 1 (p20). Western blot results from 4 experiments were quantitated using Image J and plotted below the Western images. **(E)** Macrophages were primed with Pam3 and then 5 μg/ml PGPC or POVPC encapsulated in DOTAP was added. After 18 h incubation, the media were subjected to Western blot analysis for cleaved caspase 1 (p20). The experiment was repeated with similar results. **(F)** Macrophages were treated with 1 μg/ml LPS for 21 h or primed with 1 μg/ml LPS for 3 h and then 1 μg/ml LPS mixed with DOTAP was added to deliver LPS into the cytosol to activate caspase 11. After 18 h incubation, IL-1β was measured in the culture medium. Data were combined from 2 experiments. n = 5 wells/group/experiment. **(G)** Macrophages were primed with Pam3 and then 2 mM ATP was added to induce inflammasome activation. After 15 h incubation, the culture medium was analyzed. Western blot results from 3 experiments were quantitated using Image J and plotted with the Western results. n = 5 wells/group. For IL-1β ELISA, n = 4 or 6/group.

### AOAH regulates inflammatory responses and lung injury induced by oxPLs

Having found *in vitro* that (a) rhAOAH can deacylate PGPC, POVPC, and LPC and reduce their ability to stimulate macrophages and (b) AOAH produced by resident macrophages can reduce oxPL-induced cell death and inflammasome activation, we next used four acute lung injury models to study the enzyme’s role *in vivo.* We first tested whether oxPLs can induce AOAH expression in alveolar macrophages (AMs). We found that, unlike LPS, which stimulated AOAH mRNA expression in alveolar macrophages by 100-fold *in vivo* (Zou et al., 2017), oxPLs induced AOAH expression by only 7-fold (Fig. 5A). Moreover, oxPLs reduced LPS-induced *Aoah* expression by AMs, suggesting that oxPLs may act as partial agonists to induce AOAH expression (Fig. 5A).

**Fig. 5.**
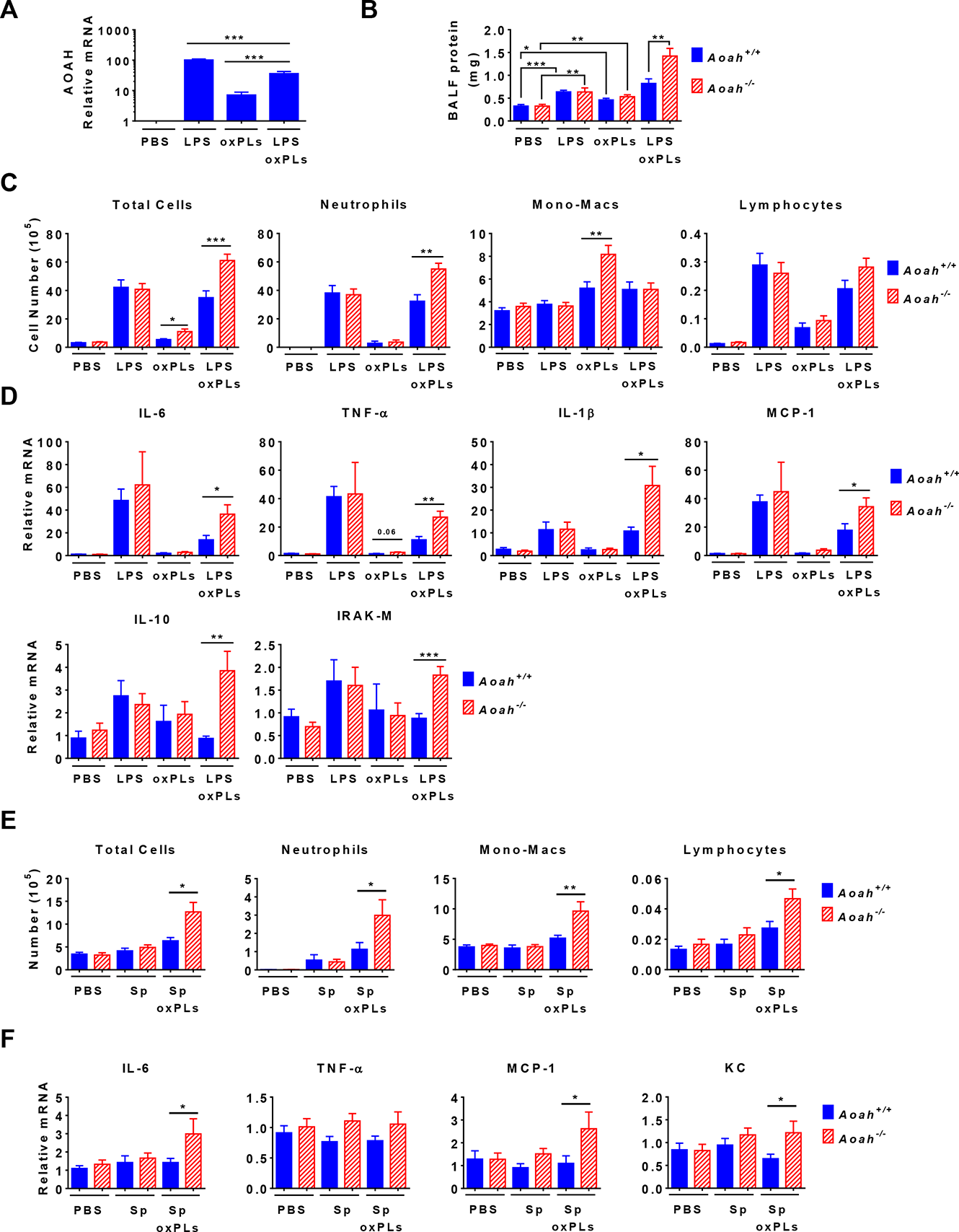
AOAH reduces inflammatory responses and lung injury induced by oxPLs. **(A - D)** *Aoah^+/+^* mice were instilled i.n. with 10 μg LPS, oxPLs (25 μg PGPC + 25 μg POVPC) or LPS + oxPLs. Eighteen h later, AOAH expression in AMs was measured using qPCR. Data were combined from 3 experiments, n = 9 mice/group (A). BALF protein amount was measured (B). BAL immune cells were analyzed (C). Lung cytokine/chemokine and IRAK-M expression were measured using qPCR (D). Data (B - D) were combined from at least 3 experiments, each with n = 3 mice/group. **(E, F)** Mice were instilled with 40 μl PBS containing 6 X 10 exp^6^ heat-inactivated *Streptococcus pneumoniae*, with or without oxPLs (25 μg PGPC and 25 μg POVPC). Eighteen h later, their BALF cells were analyzed (E). Lung cytokine/chemokine expression was measured using qPCR (F). Data were combined from at least 2 experiments, each with n = 3 mice/group.

As we had observed that sequential and simultaneous treatment of macrophages with LPS and oxPL induced similar IL-1β production (Fig. S2), we instilled *Aoah^+/+^* and *Aoah^-/-^* mice i.n. with 40 μl PBS containing 10 μg LPS, oxPLs (25 μg PGPC and 25 μg POVPC), or LPS plus oxPLs. Whereas *Aoah^+/+^* and *Aoah^-/-^* mice had similar levels of BALF protein (reflecting alveolar leakage) 18 h after either LPS or the oxPLs were introduced, *Aoah^-/-^* mice that received both LPS and oxPLs had more alveolar protein and more neutrophils in their BALF at this time than did identically treated *Aoah^+/+^* mice (Fig. 5B, C). In addition, both inflammatory (IL-6, TNF-α, IL-1β, MCP-1) and anti-inflammatory (IL- 10, IRAK-M) mRNA were more abundant in *Aoah^-/-^* mouse lungs (Fig. 5D). As AOAH can also inactivate LPS, to examine more specifically the enzyme’s role in inactivating oxPLs we also used heat-inactivated Gram-positive bacteria, *Streptococcus pneumoniae (S.p.)* to induce inflammation in the lung. We instilled heat-killed *S.p.* i.n., with or without oxPLs, to *Aoah^+/+^* and *Aoah^-/-^* mice. We did not observe significant differences in pulmonary inflammation between the two strains of mice when they received only *S.p.* instillation (Fig. 5E, F). Following instillation of *S.p.* plus oxPLs, we found that *Aoah^-/-^* mice had greater leukocyte infiltration in their airspaces and more pulmonary IL-6, MCP- 1 and KC mRNA than did *Aoah^+/+^* mice, providing further evidence that AOAH can act on oxPLs to ameliorate lung inflammation (Fig. 5E, F).

### AOAH reduces bioactive oxPLs in the lungs after HCl and oxPL instillation

In a third animal model, we used HCl and oxPLs to induce pulmonary inflammation in *Aoah^+/+^*, *Aoah^-/-^*, and transgenic mice that express high levels of AOAH in their macrophages (Ojogun et al., 2009). When we instilled only HCl and studied the mice 18 h later, *Aoah^-/-^* and *Aoah^+/+^* mice had weak and similar alveolar barrier damage (Fig 6A) and pulmonary inflammation (Fig. 6B-D). When both HCl and oxPLs were instilled, in contrast, *Aoah^-/-^* mice had much more alveolar barrier damage and lung inflammation than did *Aoah^+/+^* mice (Fig. 6A-D). In addition, the transgenic mice that express high levels of AOAH (Ojogun et al., 2009) had reduced airway inflammation after instillation of HCl plus oxPLs (Fig. S3). These findings again support a significant role for AOAH in modulating oxPL-induced pulmonary injury.

**Fig. 6.**
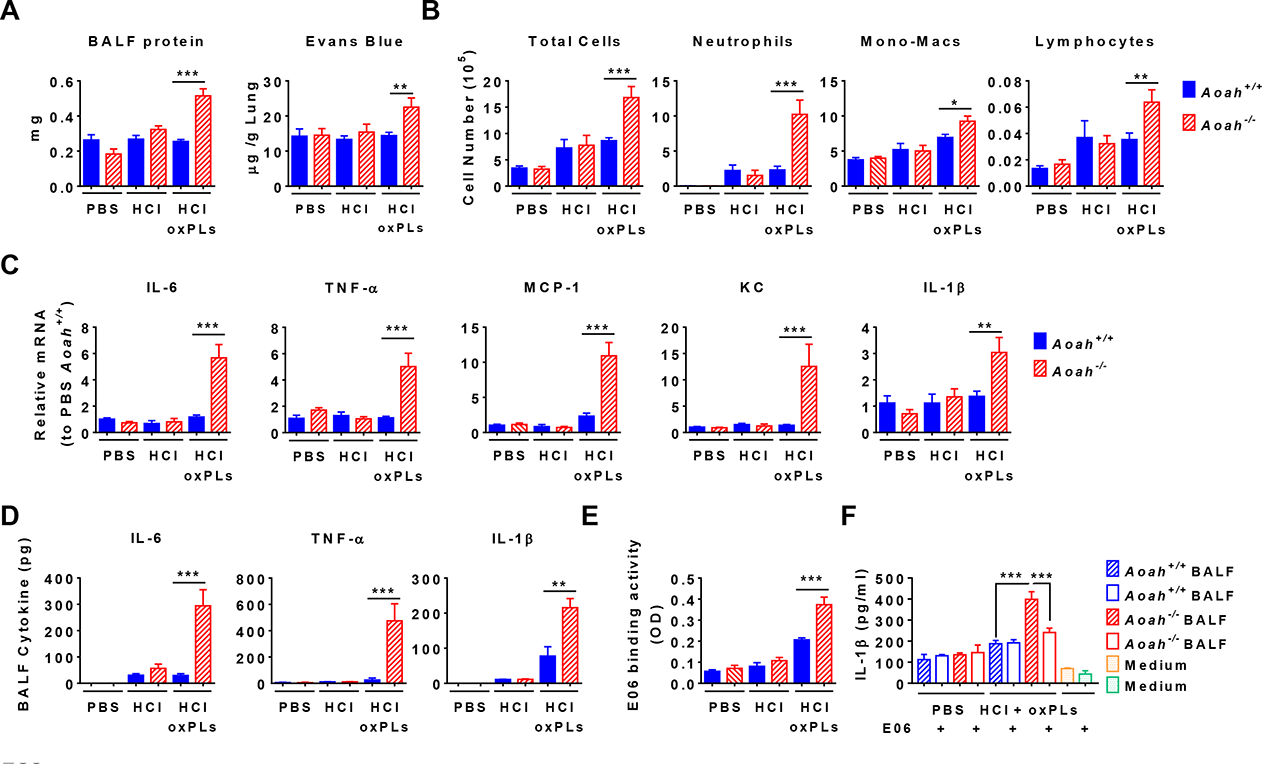
AOAH reduces bioactive oxPLs in the airspaces after HCl and oxPL instillation. **(A)** *Aoah^+/+^* and *Aoah^-/-^* mice were instilled with 40 μl 0.2 M HCl and oxPLs (25 μg PGPC + 25 μg POVPC). Eighteen h later, BALF was obtained. Protein in BALF was quantitated. Seventeen h after i.n. HCl and oxPL instillation, Evans blue was injected i.v. One hr after injection, the lung extravascular dye was extracted and measured. Data were combined from 2 or 3 experiments, each with n = 3 mice/group. **(B – F)** In the same experiments as in (A), cells in BALF were analyzed after cytospin and staining (B). Lung cytokine/chemokine expression was measured by qPCR (C). In B and C, data were combined from at least 3 experiments, each with n = 3 mice/group. Cytokines in BALF were analyzed using ELISA (D). E06 detectable oxPLs in BALF were measured (E). BALF was used to stimulate Pam3-primed macrophages for 18 h in the presence of 5 μg/ml E06 or a control mouse IgM. IL-1β in the media was measured using ELISA (F). (D - F) Data were combined from 2 experiments, each with n = 3 mice/group.

To obtain further evidence that AOAH modulates oxPL bioactivity *in vivo*, we used E06, a natural IgM that binds to the hydrophilic head of oxidized PCs such as POVPC and blocks their ability to induce inflammation *in vitro* and *in vivo* (Friedman et al., 2002; Que et al., 2018; Shaw et al., 2000; Sun et al., 2020). We detected more E06-binding activity in BALF from *Aoah^-/-^* mice, suggesting that *Aoah^- /-^* mice were less able to degrade oxPLs in their airways (Fig. 6E). We used BALF to stimulate Pam3-primed macrophages and found that BALF from HCl- and oxPL-instilled *Aoah^-/-^* mice induced significantly more IL-1β production than did BALF from identically treated *Aoah^+/+^* mice (Fig. 6F). Pre-incubation with E06 Ab diminished the ability of BALF from *Aoah^-/-^* mice to induce IL-1β production (Fig. 6F). Taken together, these results again support the conclusion that AOAH can degrade and inactivate oxPLs *in vivo*.

### AOAH regulates inflammatory responses induced by endogenous oxPLs

In the previous HCl-induced lung inflammation models, we instilled exogenous oxPLs mixed with HCl i.n. (Fig. 6). To test whether AOAH can regulate endogenous oxPL-induced lung inflammation, we used an acid aspiration-induced acute lung injury model that has been shown to induce oxPLs in the lung (Imai et al., 2008). Mice were instilled with 40 μl PBS or 0.2 M HCl, ventilated for 1 h, and allowed to recover off the ventilator for 24 h before analysis. The ventilated *Aoah^-/-^* mice had more neutrophils and more IL- 6 in their airspaces than did identically treated *Aoah^+/+^* mice (Fig. 7A, B), especially when the mice had been instilled with HCl. Ventilated *Aoah^-/-^* mice also had higher levels of lung inflammation than did the control strain (Fig. 7C). Histological study showed that HCl-instilled, ventilated *Aoah^-/-^* mouse lungs had more leukocyte infiltration and alveolar barrier thickening than did control mouse lungs (Fig. 7D). BALF from *Aoah^-/-^* mice not only bound more E06 antibody than did BALF from *Aoah^+/+^* mice (Fig. 7E), but it also had greater ability to induce IL-1β from Pam3-primed macrophages (Fig. 7F). This activity was partially inhibited by E06 (Fig 7F). These results provide additional evidence that AOAH is able to inactivate oxPLs generated *de novo in vivo*.

**Fig. 7.**
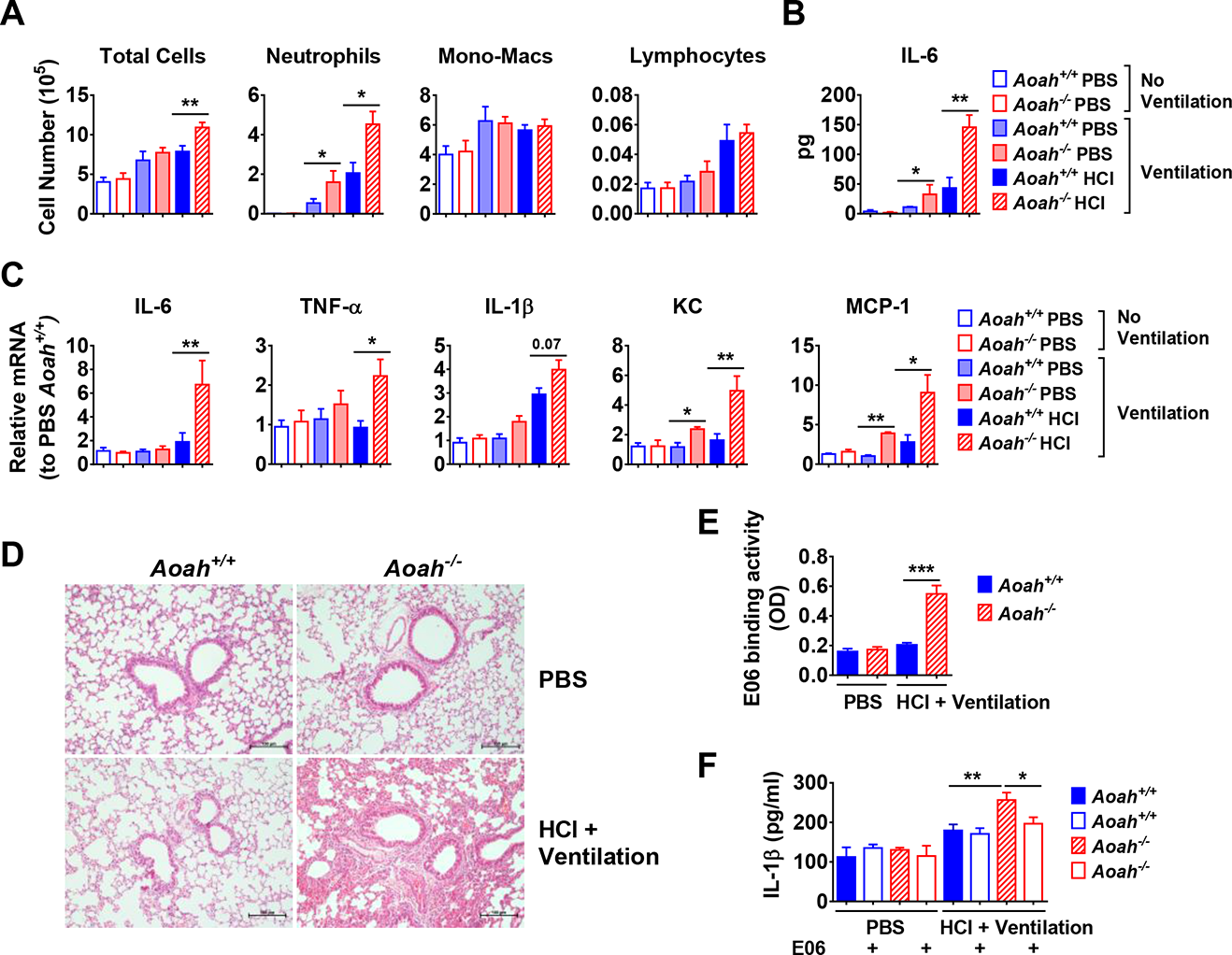
AOAH regulates inflammatory responses induced by endogenous oxPLs. *Aoah^+/+^* and *Aoah^-/-^* mice were instilled with 30 μl PBS or 0.2 M HCl, and then they were ventilated for 1 h. Twenty-four h later, the mice were analyzed. **(A)** Immune cells in BALF were counted. **(B)** IL-6 in BALF was measured using ELISA. TNF-α and IL-1β were undetectable. **(C)** Cytokine/chemokine expression was measured in the lungs. Data (A- C) were combined from at least 2 experiments, each with n = 3 mice/group. **(D)** Mouse lungs were excised and fixed in paraformaldehyde. The fixed lungs were sectioned and stained with Hematoxylin-Eosin. One representative picture of 5 in each group is shown. **(E)** BALF was collected and E06 binding activity was quantitated. **(F)** BALF was used to stimulate Pam3-primed macrophages and the IL-1β in culture medium was measured. Data were combined from 4 experiments, each with n = 3 mice/group (E and F).

## Discussion

Acting as danger-associated molecular patterns (DAMPs), oxPLs can have diverse biological activities that may contribute to the pathogenesis of acute lung injury, atherosclerosis, non-alcoholic fatty liver disease, neurodegenerative disorders and cancer (Bochkov et al., 2017; Bochkov et al., 2010). Previous studies have described several mechanisms for inactivating these phospholipids. Plasma platelet-activating factor acetyl hydrolase (PAF-AH) and intracellular PAF-AH II, which cleave sn-2 oxidized acyl chains from PGPC and POVPC, are GDSL lipases that have structural similarity to AOAH; lecithin-cholesterol-acyltransferase transfers oxidized acyl chains from oxPLs to cholesterol (Goyal et al., 1997); and oxidized acyl chains can be reduced by enzymes such as aldose reductase, phospholipid glutathione peroxidase, peroxiredoxin 6, and glutathione transferase (Mauerhofer et al., 2016; Vladykovskaya et al., 2011). Phospholipase A_2_ was implicated in POVPC deacylation in human THP-1 cells, which produce very little AOAH; the resulting LPC was more stimulatory than was POVPC itself (Vladykovskaya et al., 2011). Autophagy also helps remove oxPLs (Bochkov et al., 2010; Mauerhofer et al., 2016).

AOAH was originally found to cleave the ester bonds that attach the piggyback (secondary) fatty acyl chains to the primary 3-hydroxy acyl chains in the lipid A moiety of LPS (Hall and Munford, 1983; Munford and Hall, 1986). AOAH-deacylated (tetraacyl) LPS has greatly reduced stimulatory potency and can inhibit LPS signaling by competing with LPS for binding to LPS-binding protein, soluble and membrane CD14, and MD-2 (Erwin and Munford, 1990; Kitchens et al., 1992; Munford and Hall, 1986; Munford and Hall, 1989). Subsequent studies have found that AOAH plays a significant role in inactivating both exogenously-administered LPS and LPS derived from intestinal commensal Gram- negative bacteria (Lu et al., 2013; Lu et al., 2008; Lu et al., 2005; Qian et al., 2018; Shao et al., 2011; Shao et al., 2007; Zou et al., 2017). AOAH was also found to have phospholipase, lysophospholipase, diacylglycerollipase and acyltransferase activities *in vitro* (Gorelik et al., 2018; Munford and Hunter, 1992; Staab et al., 1994). In the present study, we found that AOAH can deacylate both oxidized glycerophospholipids and LPC and diminish their ability to induce macrophage cell death and stimulate inflammasome activation *in vitro*. Using four mouse models, we found that AOAH diminishes oxPL- induced acute lung injury.

In previous studies, we found that AOAH does not prevent acute responses to LPS. Instead, the enzyme modulates the late, or longer-lasting, responses to LPS stimulation (Lu et al., 2008; Lu et al., 2005; Shao et al., 2007; Zou et al., 2017). *Aoah^+/+^* and *Aoah^-/-^* mice had similar degrees of lung injury 24 h after LPS was instilled i.n. , while on day four, *Aoah^-/-^* mice showed delayed resolution of pulmonary inflammation (Zou et al., 2017). *Aoah^+/+^* and *Aoah^-/-^* resident peritoneal macrophages had comparable innate responses to TLR stimuli (Lu et al., 2013; Lu et al., 2008). In contrast, AOAH efficiently limits inflammasome activation by oxPLs. This may be related in part to its location in endo-lysosomes (Luchi and Munford, 1993; Staab et al., 1994), where oxPLs may be delivered when they are internalized by CD14 (Kagan et al., 2008; Zanoni et al., 2011; Zanoni et al., 2017), and by the enzyme’s ability to inactivate both oxPLs and LPCs. By inactivating oxPLs in endo-lysosomes, AOAH prevents oxPL leakage into the cytosol and endo-lysosomal damage, diminishing inflammasome activation (Fig. S4). Thus, while endo-lysosomal AOAH cannot deacylate LPS before it is sensed by TLR4, the enzyme may act on oxPLs before they activate the inflammasome.

In summary, we have presented *in vitro* and *in vivo* evidence that a host lipase, AOAH, can deacylate and inactivate two important oxPLs and also LPC. It has been intriguing that CD14 can transfer both LPS and oxPLs into endosomes (Kitchens and Munford, 1998; Zanoni et al., 2017) and that TLR4 and caspase 11 can recognize both oxPLs and LPS (Hagar et al., 2013; Imai et al., 2008; Kayagaki et al., 2011; Kayagaki et al., 2013; Shi et al., 2014; Zanoni et al., 2016); here we show that the same lipase can inactivate both of these agonists and prevent them from inducing inflammasome activation. As oxPLs and lysophospholipids can contribute to many inflammatory diseases and aging, inactivating these lipids may be a very important function of this highly conserved enzyme.

## Materials and Methods

### Ethics statement

C57BL/6J *Aoah^+/+^*, *Aoah^-/-^*, and transgenic CD68-AOAH mice were produced at the University of Texas Southwestern Medical Center, Dallas, Texas, and provided to the National Institute of Allergy and Infectious Diseases, Bethesda, MD, and Fudan University, Shanghai (Lu et al., 2003; Lu et al., 2005). Male and female mice 6 - 8 weeks old were used. All mice were studied using protocols approved by the Institutional Animal Care and Use Committee (IACUC) of Fudan University and the National Institute of Allergy and Infectious Diseases. All protocols adhered to the Guide for the Care and Use of Laboratory Animals.

### Prime and stimulate macrophages

Peritoneal cells were harvested from naïve mice and cultured in RPMI medium (Gibco) containing 10% fetal bovine serum (Gibco), 2 mM glutamine, 100 U/ml penicillin, and 0.1 mg/ml streptomycin (Life Technologies). After incubation for 4 h at 37℃ to let the macrophages adhere to culture plates, the floating cells were washed away and the peritoneal macrophages were incubated in RPMI medium containing 0.1% FBS, 100 U/ml penicillin, and 0.1 mg/ml streptomycin with or without 10 ng/ml Pam3CSK4 (tlrl-pms, InvivoGen), 10 ng/ml LPS O111:B4 (Sigma) for 7 h to prime macrophages. 1- palmitoyl-2-glutaryl-sn-glycero-3-phosphocholine (PGPC, Cayman), 1-palmitoyl-2-(5’- oxo-valeroyl)-sn-glycero-3-phosphocholine (POVPC, Cayman), 1-palmitoyl-2- arachidonoyl-sn-glycero-3-phosphocholine (PAPC, Sigma), oxPAPC (oxidized PAPC, Avanti) or 1-palmitoyl-2-hydroxy-sn-glycero-3-phosphocholine (LPC, Cayman) in ethanol were dried under nitrogen and then suspended in RPMI containing 0.1% FBS with vigorous vortexing, water bath sonication, and further vortexing. The lipids were then added to culture medium to stimulate primed or unprimed macrophages for 18 h. We used phospholipid concentrations that were below their critical micellar concentrations, which are 54 μM, 68 μM, and 50 μM for PGPC, POVPC and LPC respectively, and similar to levels found in animal tissues (Bochkov et al., 2010; Fruhwirth and Hermetter, 2008; Mauerhofer et al., 2016; Pande et al., 2010). Brief incubation in 10% serum, used for many *in vitro* studies of oxPL activities, has been reported to convert POVPC and PGPC to LPC (Stemmer et al., 2012). We used 0.1% serum to avoid this issue.

We used DOTAP (N-[1-(2,3-Dioleoyloxy)propyl]-N,N,N-trimethylammonium methyl-sulfate) (Roche) to introduce LPS or oxPLs into cells following the method by Zanoni et al (Zanoni et al., 2016). Briefly, 375 ng DOTAP was used to encapsulate 5 μg LPS or 10 μg oxPLs in 10 μl Opi-MEM (Thermo). After 30 min incubation at 37℃, the DOTAP/LPS or DOTAP/oxPLs were diluted and added to cell culture at final concentrations of 1 μg/ml or 5 μg/ml respectively.

### Cell death analysis

Crystal violet (CV) (Sigma) was used to measure adherent cell DNA as described (Feoktistova et al., 2016). The culture medium was used for the LDH cytotoxicity assay (Pierce) according to manufacturer’s instruction.

### Western blot and ELISA

After macrophages were primed with Pam3 and challenged with oxPLs, cell-free medium was collected and medium protein was precipitated using the chloroform-methanol method of Fic et al. (Fic et al., 2010). The protein pellet was dissolved in SDS loading buffer containing 10% β-mercaptoethanol and subjected to Western blotting analysis. IL-1β and caspase 1 p20 were detected using monoclonal antibodies D4T2D and E2G2I respectively (Cell Signaling Technology). Mouse IL-1β and IL-6 ELISAs (BD) were performed according to the manufacturer’s instructions.

### AOAH activity toward PGPC, POVPC, and LPC *in vitro*

PGPC and POVPC (Cayman) (100 nmoles) were dried under argon before 5 μl ethanol was added to facilitate resuspension. Fatty acid-free BSA in 0.9% saline (0.2 mg/ml final concentration) and Triton X-100 (0.09%) were added while vortexing under argon and the suspension was briefly sonicated in a water-bath. Ten mM sodium acetate was added to adjust pH to 5.9 - 6.1. After mixing thoroughly and removing 50 μl (no AOAH control), 1 μl of 1 mg/ml recombinant human acyloxyacyl hydrolase (rhAOAH, provided by ZymoGenetics and stored at -80℃ (Hagen et al., 1991)) was added to 450 μl reaction mixture, 50 μl of the mixture were aliquoted into polypropylene microfuge tubes, and incubation was carried out at 37℃ with intermittent rotation for 2 or 4 h. Ten μl were removed from each tube to measure C8 or longer free fatty acids (i.e., palmitic acid released from PGPC or POVPC) (Sigma free fatty acid kit MAK044) and the remaining samples were stored at -80℃ prior to use or analysis.

The AOAH-treated PGPC and POVPC reaction products were analyzed using an Agilent 1290 Infinity UPLC coupled to a 6460C Agilent triple quadrupole mass selective detector (LC-QqQ). Samples were separated on an EclipsePlus C18 column 2.1 mm x 50 mm 1.8 mm particle, using aq. 0.1% formic acid (mobile phase A) and acetonitrile with 0.1% formic acid (mobile phase B). Flow rate was 0.8 mL/min beginning with 5% B holding 0.25 min, ramping to 95% B over 9 min, hold 4 min, and re-equilibrate. Mass Spectrometer was operated in multiple reaction monitoring (MRM) mode with M+H^+^ion as the precursor and product ion m/z 184.1 (phosphatidyl choline group) created using CEV (collision energy voltage) 24 - 28 V. Fractions were calculated using % signal abundance observed in MRM.

To treat LPC, 1 μg rhAOAH was added to 450 nmoles LPC as above, and the pH was adjusted with 10 mM Na acetate to pH 5.9 - 6.1. After 2 or 4 h incubation at 37℃, the released palmitate was measured using the Sigma free fatty acid assay kit, which measures fatty acids with ≥ 8 carbons. No free fatty acid was detected when AOAH was not added.

### Caspase inhibitors

Ten μM VX-765 (caspase-1 inhibitor, Selleck) or 20 μM Z-VAD-FMK (pan-caspase inhibitor, Selleck) was added to the culture medium for 1 h before macrophages underwent the processes of priming and stimulation as described previously.

### Animal models

We used four lung injury models: LPS-induced, gram-positive bacterial infection (*Streptococcus pneumoniae*)-induced, acid gastroesophageal reflux (HCl), and acid aspiration-ventilation-induced. Mice were randomly assigned to different treatment or control groups.

*Streptococcus pneumoniae bacteria* (*Sp*) were cultured on Columbia blood agar plates and a single colony was picked and inoculated in 3% TSB (Tryptic Soy Broth) medium containing 10% FBS. After shaking at 150 rpm, 37℃ for 10 h, bacteria were collected by centrifugation, re-suspended in PBS and then heat-inactivated at 100℃ for 10 min. The bacterial culture with 6.6 x 10^8^ *Sp*/ml contained 0.012 EU/ml endotoxin, determined using the Endotoxin Detection kit (InvivoGen).

#### 1) Three intranasal instillation acute lung injury models

Fifty μg OxPLs (25 μg PGPC + 25 μg POVPC) in ethanol were added to a polypropylene tube and the ethanol was evaporated using nitrogen. (a) Ten μg LPS, (b) 6 × 10^6^ heat-inactivated *Streptococcus pneumoniae* bacteria suspended in 40 μl PBS or (c) 40 μl 0.2 M HCl was used to re-suspend dried 50 μg oxPLs. Each of the suspensions then underwent vortexing for 30 sec, water bath sonication for 15 min, and vortexing for another 30 sec before it was instilled intranasally into mice which had been anesthetized i.p. with 0.5% pentobarbital sodium (50 μg/g body weight). After 18 h, mice were exsanguinated by cutting the inferior vena cava. Bronchoalveolar lavage

(BAL) was performed by cannulating the trachea with a 20-gauge catheter that was firmly fixed with a suture. The lung was then infused with 1 ml PBS containing 5mM EDTA and bronchoalveolar lavage fluid (BALF) was harvested. This procedure was repeated 5 times and the BALF was combined. The lung was then cut into pieces for preservation in RNA Later (TianGen) for RT-PCR or in 4% paraformaldehyde for histological analysis.

In some experiments we used transgenic mice that express AOAH from the CD68 promoter in macrophages and dendritic cells (Ojogun et al., 2009).

#### 2) Acid aspiration-ventilation acute lung injury model

Mice were anesthetized i.p. with 1% pentobarbital sodium (100 μg/g body weight) to reach a state of deep anesthesia. An infrared heater was used to maintain body temperature. Forty μl 0.2 M HCl was instilled intratracheally using a 1 mm diameter capillary tube. A 20-gauge catheter was then inserted into the airway and connected to a mechanical respirator (Harvard apparatus). Respiration was carried out for 60 min with 100 breaths/min, tidal volume 0.5 ml, sigh volume 0.5 ml, sigh frequency 1/0 and inspiratory to expiratory ratio 1:2. The mice were then removed from the respirator and monitored. Twenty-four h later, mice were euthanized and analyzed.

### Analysis of bronchoalveolar lavage fluid

After the lungs were lavaged with 1 ml PBS twice, BALF was collected and centrifuged. The cell-free supernatant was used to measure protein concentrations with a bicinchoninic acid (BCA) kit (Pierce). The total BALF protein amount equals protein concentration times 1 ml. Inflammatory cytokines and chemokines in BALF were measured using TNF-α, IL-6, or IL-1β ELISA kits (BD). The cell pellet was re- suspended in PBS and total cell numbers were counted using Cellometer (Nexelcom). Cell differential counting was conducted using cytospin and Wright-Giemsa staining. After counting 300 cells, the proportions of neutrophils, monocytes-macrophages and lymphocytes were calculated and used to determine the number of each cell type in the BALF. To assess alveolar barrier damage, extravascular Evans blue were measured as described previously (Zou et al., 2017). Briefly, Evans Blue was injected i.v. 1 h before euthanasia. After perfusion of the lungs, extravascular Evans blue was extracted and the optical density was measured at 620 nm (reference 740 nm) using a Tecan reader, as described previously (Zou et al., 2017).

### Real time-PCR

RNA from AMs or lungs was isolated using RNA isolation kit (Tiangen) and reversely transcribed (Tiangen). The following primers were used at a concentration of 10 μM: ao ah (5’-CAGCTACTCCCATGGCCAAA-3’, 3’-GCCACCTGGACTGAAGAGTT-5’), actin (5’-GGCTG TATTCCCCTCCATCG-3’, 3’-CCAGTTGGTAACAATGCCATGT-5’), il-6 (5’-ATCGTGGAAATGAG AAAAGAGTTGT-3’, 3’-AAGTGCATCATCGTTGTTCATACA-5’), tnf-α (5’-CATCTTCTCAAAA TTCGAGTGACAA-3’, 3’-TCAGCCACTCCAGCTGCTC-5’), mcp-1 (5’-TCCCAATGAGTAGGCTGG AG-3’, 3’-TCTGGACCCATTCCTTCTTG-5’), il-1β (5’-CCTTCCAGGATGAGGACATGA-3’, 3’-GTC ACACACCAGCA GGTTATCA-5’), kc (5’-CAAGAACATCCAGAGCTTGAAGGT-3’, 3’-GTGGCTAT GACTTCGGTTTGG-5’), il-10 (5’-GCTGGACAACATACTGCTAACC-3’, 3’-ATTTCCGATAAGGCT TGGCAA-5’), irak-m (5’-TCCCACCTGAGGTGAAGCAT-3’, 3’-TGTGACATTGGCTGGTTCCA-5’). Actin was used as the internal control and relative gene expression was calculated by t he ΔΔCt quantification method.

### Histology

Lungs were excised and fixed in 4% paraformaldehyde containing 5% sucrose and 2 mM EDTA for 24 h. The lung tissues were sectioned and stained with hematoxylin and eosin (H&E). The samples were examined using a Nikon E200 microscope blinded.

### Detection of BALF oxidized PCs by E06 Ab

A published method was used with modification (Imai et al., 2008). Briefly, after BALF was harvested and centrifuged, 100 μl supernatant was transferred to a high binding polystyrene plate (Corning) and incubated at 4℃ for 18 h. The plate was washed 3 times with PBS and blocked with 5% BSA in Tris-Buffered Saline pH 7 for 1 h. Five μg/ml E06 antibody (Avanti Polar Lipids) or a control mouse IgM (clone, MM-30, BioLegend) was added and incubated at room temperature for 2 h. After washing the plate 3 times with PBS, anti-mouse IgM (H+L)-HRP antibody (Sigma) in 100 μl PBS was added and incubated at room temperature for 1 h. The wells were washed 7 times with PBS and TMB substrate (BD) was added (100 μl/well). After 30 min incubation at room temperature, the plate was read at 450 nm (reference 570 nm) (Tecan). OxPL-binding activity (OD) = OD with E06 – OD with the control IgM.

### OxPL neutralization by E06 Ab

In the lung inflammation models, the lungs were lavaged with 1 ml PBS twice and the BALF was centrifuged. The supernatant was transferred into a new tube and then incubated with 5 μg/ml E06 or a control mouse IgM for 30 min at 37℃. Pam3-primed macrophages were incubated with the BALF at 37℃ for 18 h and medium IL-1β was measured.

### Statistical analysis

Independent biological replicates are presented as mean ± SEM. Unless indicated otherwise, differences between groups were assessed using the Mann-Whitney nonparametric test (two-tailed, unpaired). *, P < 0.05; **, P < 0.01; ***, P < 0.001.

## Acknowledgments

This study was supported by grants 31770993, 91742104 and 31570910 from National Natural Science Foundation of China (M.L.), grant 21ZR1405400 from Shanghai Committee of Science and Technology (M.L.) and by the Divisions of Intramural Research, National Institute for Allergy and Infectious Diseases, NIH, USA (R.S.M, M.G.). We thank Feng Shao for providing *Caspase11^-/-^* mice.

## Supplemental figures and legends

**Fig. S1.**
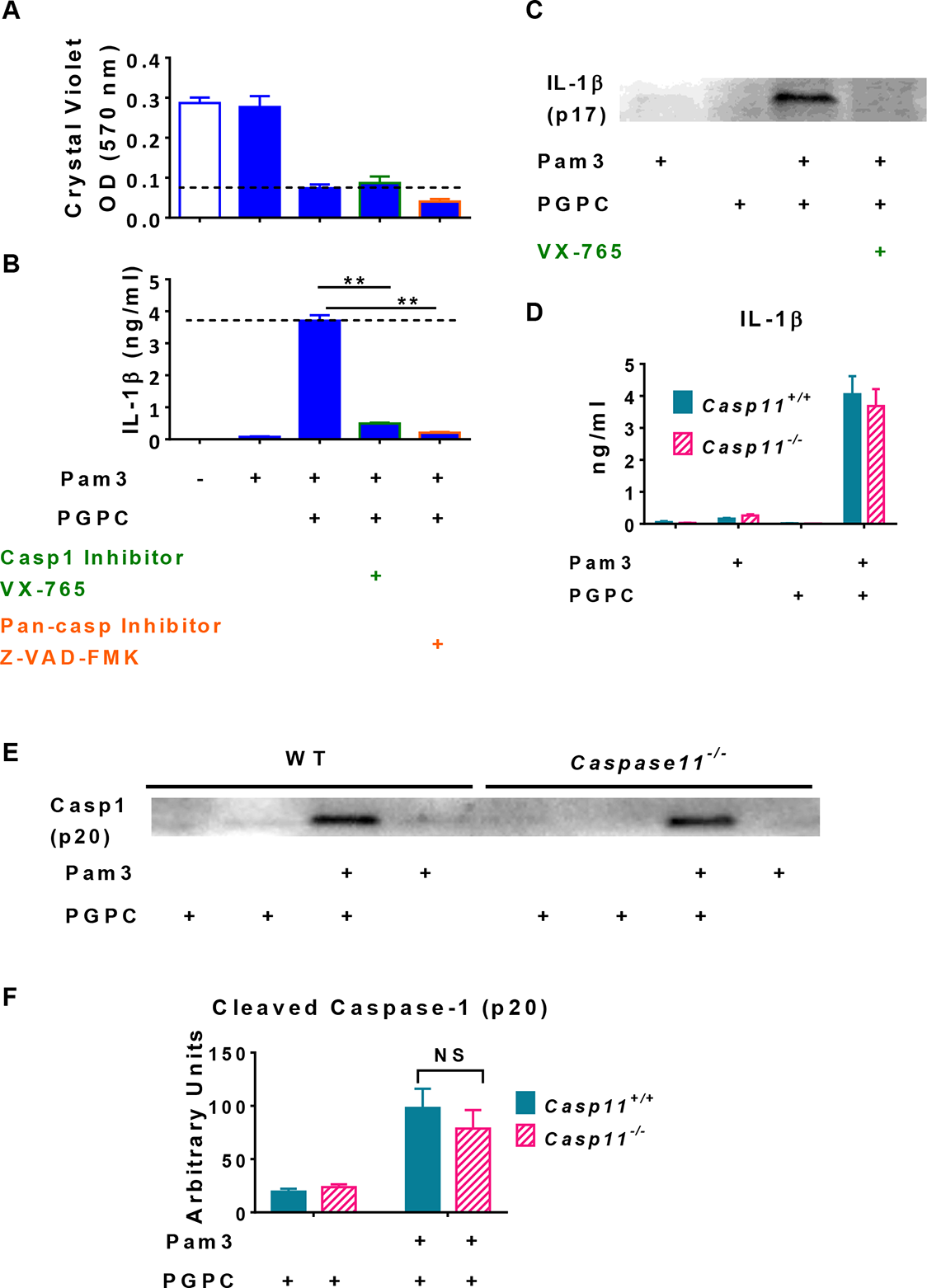
Inflammasome activation is not required for oxPL-induced macrophage death; caspase 11 is not required for PGPC-induced IL-1β release in peritoneal macrophages. **(A - C)** Peritoneal macrophages were treated with 10 μM caspase 1 inhibitor VX-765 or 20 μM pan-caspase inhibitor Z-VAD-FMK for 1 h before macrophages were primed with 10 ng/ml Pam3 for 7 h. PGPC (12.5 μg/ml) was then added. After 18 h incubation, cells were stained with crystal violet (A); the culture media were used for IL-1β ELISA **(A)** and Western blot analysis (C). Data were combined from 2 experiments, n = 6. **(D - F)** *Caspase 11^+/+^* or *caspase 11^-/-^* peritoneal macrophages were treated with 10 ng/ml Pam3 for 7 h, and then 12.5 μg/ml PGPC was added. After 18 h incubation, IL- 1β in culture medium was determined. Data were combined from 3 experiments, n= 3 or 4/group/experiment (D). Cleaved caspase 1 (p20) in the culture medium was measured by Western Blot (E); results from 3 experiments were quantitated using Image J (F).

**Fig. S2.**
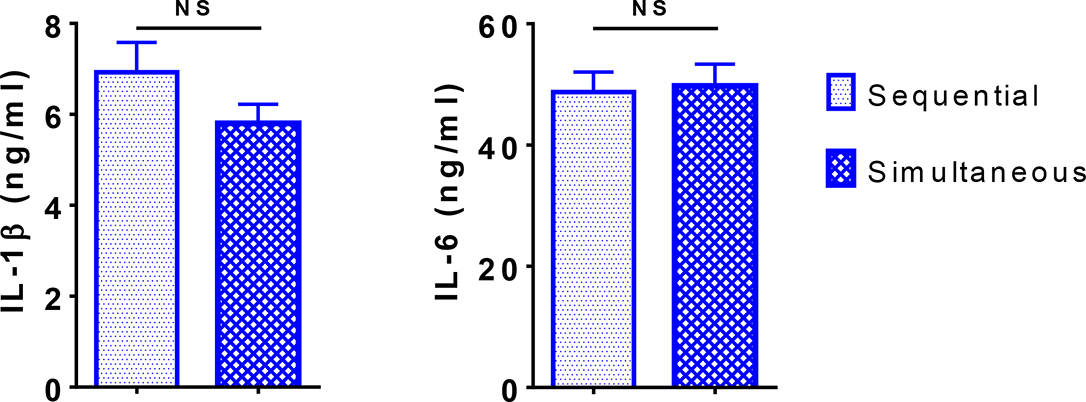
Sequential and simultaneous treatment of macrophages with LPS and oxPL induced similar IL-1β production. Peritoneal macrophages were treated with 10 ng/ml LPS for 7 h followed by treatment with 5 μg/ml PGPC for 18 h (sequential) or 10 ng/ml LPS mixed with 5 μg/ml PGPC for 18 h (simultaneous). Medium IL-1β and IL-6 were measured by using ELISA. Data were combined from 2 experiments, n = 8 or 13.

**Fig. S3.**
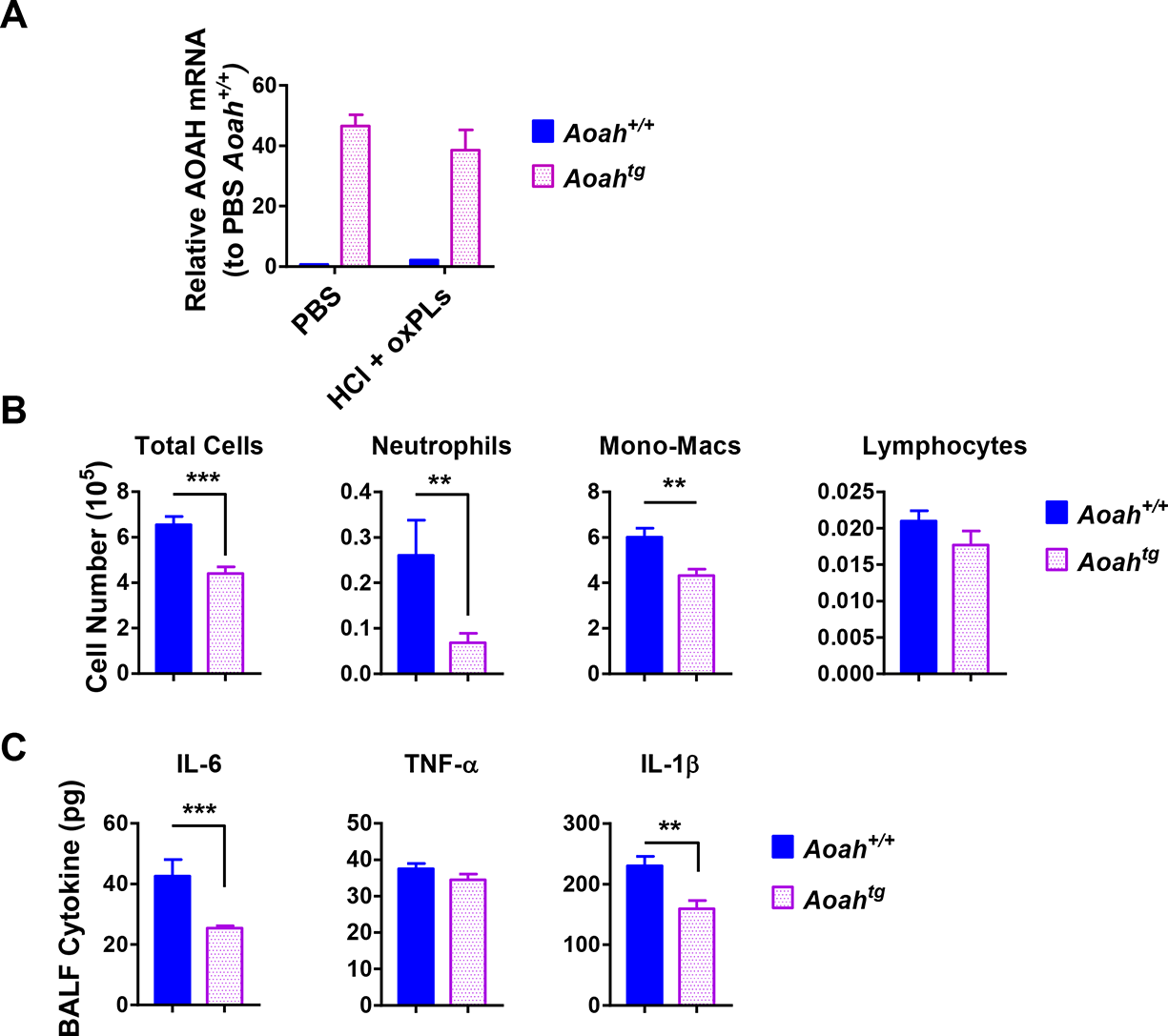
Mice that overexpress AOAH in macrophages have reduced airway inflammation after HCl and oxPL challenge. *Aoah^+/+^* and *Aoah* transgenic mice were instilled with 40 μl 0.2 M HCl and oxPLs (25 μg PGPC + 25 μg POVPC). Eighteen h later, BALF was obtained. **(A)** AOAH mRNA in the lungs of *Aoah^+/+^* and *Aoah^Tg^* mice was measured. n = 5/group. **(B)** Immune cells in BALF were analyzed after cytospin and staining. **(C)** BALF cytokines were measured using ELISA. (B and C) Data were combined from 2 experiments, n = 9-15/group.

**Fig. S4.**
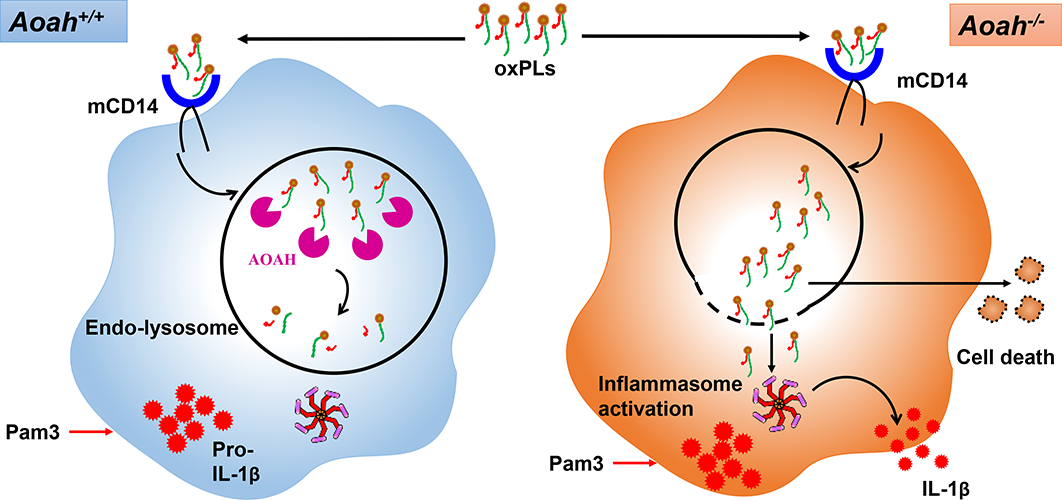
AOAH deacylates and inactivates oxPLs. After oxPLs enter endo-lysosomes in a CD14-dependent pathway, AOAH in endo- lysosomes deacylates and inactivates oxPLs. When AOAH is lacking, oxPLs leak into cytosol or endo-lysosome membrane ruptures, leading to inflammasome activation.

